# Dietary protein increases T cell independent sIgA production through changes in gut microbiota-derived extracellular vesicles

**DOI:** 10.1101/2020.11.30.405217

**Authors:** Jian Tan, Duan Ni, Jemma Taitz, Gabriela Veronica Pinget, Mark Read, Alistair Senior, Jibran Abdul Wali, Ralph Nanan, Nicholas Jonathan Cole King, Georges Emile Grau, Stephen J. Simpson, Laurence Macia

**Affiliations:** Charles Perkins Centre, The University of Sydney, Sydney, NSW, Australia; School of Medical Sciences, Faculty of Medicine and Health, University of Sydney, Sydney, NSW, Australia; School of Computer Science, Faculty of Engineering, University of Sydney, Sydney, NSW, Australia; School of Life and Environmental Sciences, University of Sydney, Sydney, NSW, Australia; Sydney Medical School Nepean, University of Sydney, Sydney, NSW, Australia; Sydney Cytometry, The University of Sydney and The Centenary Institute, Sydney, NSW, Australia; Vascular Immunology Unit, The University of Sydney, Sydney, NSW, Australia

## Abstract

Secretory IgA (sIgA) is a key mucosal component ensuring host-microbiota mutualism. Using nutritional geometry modelling in mice fed 10 different macronutrient-defined, isocaloric diets, we identified dietary protein as the major driver of sIgA production. Protein-driven sIgA induction was not mediated by T cell-dependent pathways or changes in gut microbiota composition. Instead, the microbiota of high-protein fed mice produced significantly higher quantities of extracellular vesicles (EV), compared to those of mice fed high-carbohydrate or high-fat diets. These EV activated TLR4 to increase the epithelial expression of IgA-inducing cytokine, APRIL, B cell chemokine, CCL28, and the IgA transporter, PIGR. We showed that succinate, produced in high concentrations by microbiota of high-protein fed animals, increased the generation of reactive oxygen species by bacteria, which in turn promoted EV production. This is the first report establishing a causal link between dietary macronutrient composition, gut microbial EV release and host sIgA response.

## Introduction

The gut microbiota, defined by the trillions of bacteria that inhabit the gut, play a critical role on the host’s physiology and immunity^1^. To maintain a symbiotic relationship, strategies have been developed by the host to regulate its interaction with the gut microbiota. Mucosal secretory IgA (sIgA) play a key role in this host-microbiota mutualism by excluding pathobionts and pathogens and limiting bacterial attachment to the epithelium. While patients with selective IgA deficiency appear healthy, they are more susceptible to various diseases, including gastrointestinal disorders and allergies^2^, highlighting the importance of IgA in mucosal homeostasis.

Under homeostatic conditions, IgA is primarily produced by plasma cells local to the small intestine lamina propria through a T cell-independent pathway. The majority of IgA produced in the gut is in dimeric form, which can be transported through the epithelium into the lumen by binding to polymeric immunoglobulin receptor (pIgR)^3^. The resulting sIgA generated via this pathway is of low affinity and is polyreactive towards common bacterial antigens, limiting the attachment of diverse bacteria to the host epithelium^4^. The induction of IgA production is regulated by the local cytokine environment, with the increase in class switch recombination (CSR) cytokines, APRIL and BAFF, produced by epithelial cells, promoting the differentiation of IgM^+^ B cells into IgA-producing plasma cells. Similarly, pIgR expression can be upregulated by several inflammatory cytokines such as IFN-γ, IL-1, IL-17, TNF, and IL-4^3^. Of note, IL-4 can have a dual role by also promoting IgA CSR. TLR activation by commensal bacteria is critical both for the recruitment of IgM^+^ B cells in the lamina propria, by increasing the expression of the gut homing chemokine CCL28^5^, and for the induction IgA CSR by upregulating APRIL^6^. TLR4 is the major TLR expressed on the epithelium and its overstimulation in TLR4 transgenic mice has been shown to promote CCL28 and APRIL production and accumulation of IgA in the lamina propria and gut lumen^5^. Seminal work utilizing germ-free mice has established that the presence of gut bacteria is the main driver of sIgA response, and transient colonization of germ-free animals with *E. coli* leads to transient sIgA production^7^. However, how dynamic changes to the gut microbiota composition affect sIgA is less clear. Diet composition is a major driver of the microbiota composition with most studies focusing on microbiota profiling in the cecum and the colon^8^, where IgA production is minimal, compared to the small intestine. How dietary macronutrient composition (protein, fat and carbohydrate) affects the dynamic interplay between the small intestine microbiota and mucosal IgA remains unknown. This knowledge could establish which diet compositions are beneficial in restoring gut homeostasis and which diets are detrimental.

In our approach, we fed mice on one of 10 different isocaloric diets varying in their macronutrient composition and quantified sIgA. By using mixture modelling, we identified that dietary protein dramatically increased luminal sIgA levels, which is associated with increased expression of CCL28 and APRIL in the small intestine. These changes were correlated with the increased capacity of the microbiota to stimulate TLR4 directly or indirectly via increased production of gut microbiota-derived extracellular vesicles (EV). We also established that EV derived from high-protein diet microbiota could directly promote the expression of CCL28. Our work highlights the key role of dietary protein on sIgA production and identifies bacterial-derived EV as a novel mediator of gut microbiota-host mutualism.

## Results

### High protein feeding promotes high lamina propria IgA production and higher luminal sIgA

To determine how dietary macronutrients might affect sIgA and thus host-microbiota mutualism, we fed mice on one of 10 isocaloric diets with defined ratios of macronutrients in the ranges 5-60% protein, 20-75% fat and 20-75% carbohydrate, for 5 weeks (Fig. 1a and Supplementary Table 1).

**Figure 1:**
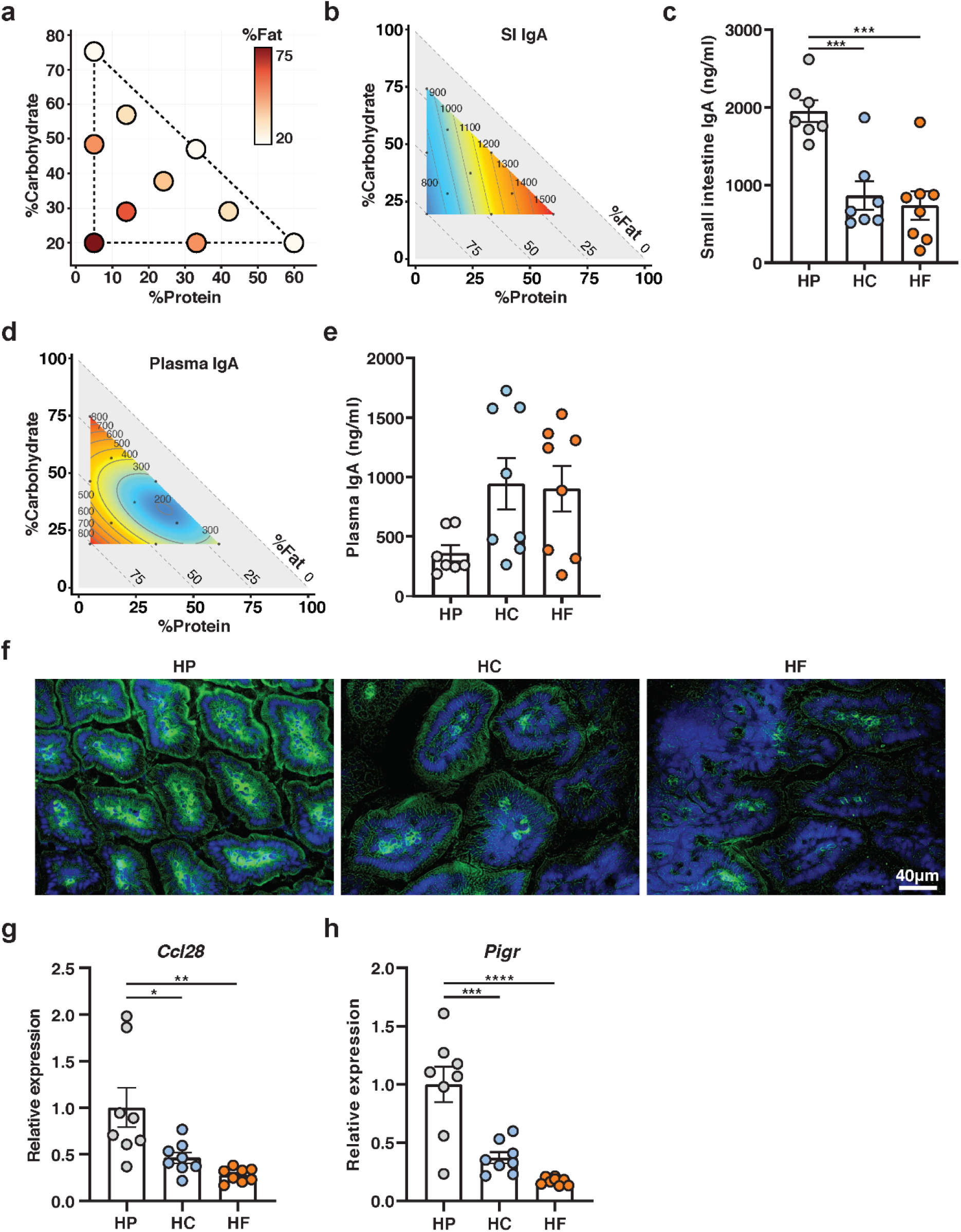
High protein feeding promotes high lamina propria IgA production and higher luminal sIgA. (a) Visual representation of the composition of the diets used in this study. Each diet is represented by one circle each and their localisation on the x-axis and on the y-axis define their proportion of protein and of fat, respectively. The proportion of carbohydrate is indicated by the colour range as illustrated in the legend. (b) Contribution of macronutrient composition to small intestinal luminal sIgA (n=7-8 per diet, quantified by ELISA) was modelled by mixture model and represented on a right-angled mixture triangle comprising of fat (y-axis), protein (x-axis) and carbohydrate (hypotenuse) with small intestinal content IgA concentration (ng/ml, numbers on isolines) as the response variable. Red represents high levels of sIgA while blue represents low levels of sIgA in the nutrient mixture space. Each dot represents one of the 10 diets used for modelling response surface. (c) Scatter bar plot of sIgA from mice fed on a HP, HC or HF diet as determined by ELISA (n=7-8 per group). (d) Mixture model of plasma IgA represented on a right-angled mixture triangle and (e) corresponding scatter bar plot. (f) Representative immunofluorescence staining of IgA (green) in the small intestine counterstained with DAPI (blue) from mice fed on a HP, HC, or HF diet for 5 weeks. Scale bar represents 40µm (h-i) Gene expression of (g) Ccl28 and (h) Pigr in whole small intestine tissue was determined by qPCR from mice fed on either a HP, HC, or HF diet (n=8 per group). Data are represented as mean ± SEM.

The impact of macronutrient composition on gut luminal concentrations of sIgA was visualised using a proportion-based nutritional geometry approach, as described previously^11,12^. Mixture models were used to determine the effects of diet composition on luminal sIgA concentration, as quantified by ELISA (n=8 per diet). Predicted effects of diet on sIgA were mapped onto a right-angled mixture triangle plot, where protein concentration in the diet is represented on the x-axis, fat on the y-axis and carbohydrate on the hypotenuse (Fig. 1b). Regions of the nutrient mixture space appearing in red demonstrate high levels of sIgA while areas in deep blue represent low levels of sIgA, and values on the isolines indicate the modelled concentration of sIgA (ng/ml). For sIgA, model 1 had the most favourable AIC (Supplementary Table 2), which reveal that dietary protein concentration was the principal predictor of sIgA, there being a clearly graded increase in sIgA with increasing protein concentration, irrespective of fat or carbohydrate content. Linear regression revealed that this effect was driven significantly by protein, but not by carbohydrate or fat intake, and was independent of total caloric intake (Supplementary Fig. 1a-d). Accordingly, when the three diets located at the apices of the mixture space in Fig. 1a were compared, the highest concentrations of gut luminal sIgA were observed in mice fed on the highest protein diet (HP; P60 C20 F20), compared to those fed on a diet high in fat (HF; P5 C20 F75) or high in carbohydrate (HC; P5 C75 F20) (Fig. 1c).

The shape of the response surface for plasma IgA was markedly different to that of sIgA, with plasma IgA being lowest on diets containing high protein coupled with both low fat and low carbohydrate (Fig. 1d-e and Supplementary Table 3). Macronutrient composition was not a predictor of plasma IgM concentrations, as the null model was determined to be most favourable by AIC (Supplementary Table 4). This suggests that high luminal sIgA under high protein feeding condition was not linked to a systemic increased in basal B cell activity and that this effect was, therefore, mucosal-specific.

To determine whether elevated sIgA levels in the gut lumen of HP fed mice was due to higher local production of IgA in the lamina propria, we performed immunofluorescence staining with anti-IgA on frozen sections of small intestine isolated from mice fed on HP, HC or HP diets. Immunofluorescence analysis showed that HP feeding led to the highest expression of IgA in the lamina propria compared to HC or HF feeding (Fig. 1f). Prior to differentiation into IgA- producing plasma B cells, B cells are recruited to the gut by the gut epithelial chemokine CCL28. By qPCR analysis, we found that mice fed on HP diet had significantly higher intestinal gene expression of the B cell gut homing chemokine, CCL28 (Fig. 1g). Additionally, the expression of the gene encoding for pIgR, involved in the transport of sIgA to the lumen via the epithelium, was significantly increased under HP feeding conditions (Fig. 1h).

Altogether, these data show that protein is the major macronutrient driving sIgA production in the gut lumen. A HP diet promotes the expression of the B cell gut homing chemokine, CCL28, and accordingly, higher presence of IgA in the small intestine lamina propria. Finally, a HP diet also promotes increased expression of pIgR, the transporter of IgA in the gut lumen, consistent with the highest concentration of luminal sIgA.

### High protein feeding does not promote IgA production through T-cell dependent mechanisms

IgA-producing plasma cells in the lamina propria either originate from IgM-producing B cells that differentiate locally in the lamina propria, or from IgA-expressing plasmablasts induced in gut-associated lymphoid structures such as the Peyer’s patches or the mesenteric lymph nodes, which migrated back towards the lamina propria to differentiate into mature plasma cells. To identify the origin of the IgA^+^ plasma cells that are increased under HP feeding condition, we assessed by flow cytometry the proportion of total B cells, as well as IgA plasmablasts in both Peyer’s patches, the main site of T-cell dependent IgA plasma cell induction, and mesenteric lymph nodes of mice fed on HP, HC or HF diets. We found that proportions of total B cells in both the Peyer’s patches (Fig. 2a) and mesenteric lymph nodes (Fig. 2b) were similar between groups. Likewise, no changes in the proportion of IgA^+^B220^+^ plasmablast were observed between groups in both the Peyer’s patches (Fig. 2c-d) and the mesenteric lymph nodes (Supplementary Fig. 2a). This suggests that HP feeding does not increase T-cell dependent IgA production, as confirmed by the similar proportions of GL7^+^CD95^+^ germinal centre B cells in both the Peyer’s patches (Fig. 2e-f) and the mesenteric lymph nodes (Supplementary Fig. 2b). Our results suggest that HP feeding promotes high levels of small intestine sIgA via a T cell-independent pathway.

**Figure 2:**
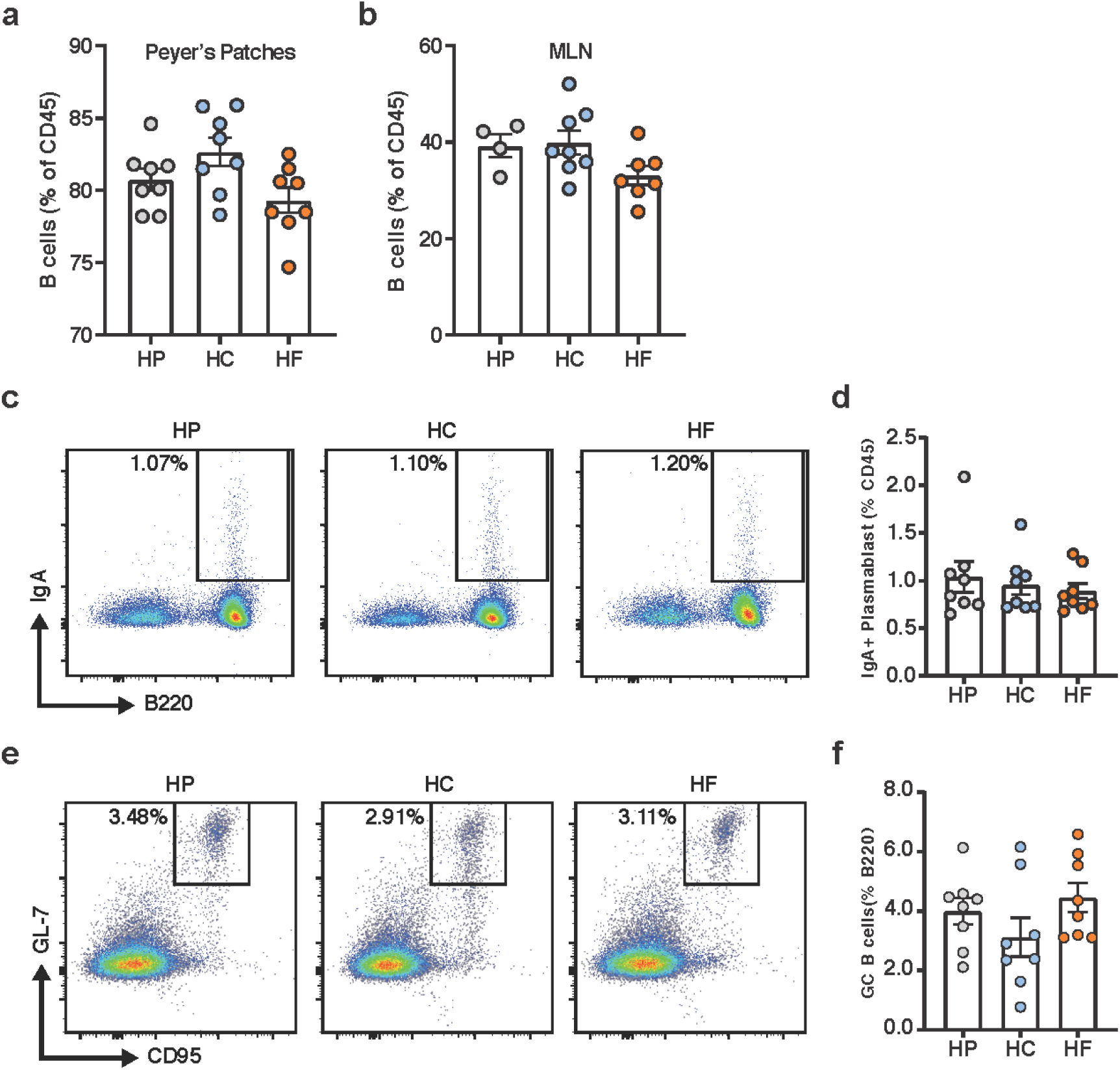
High protein feeding does not promote IgA production through T cell-dependent mechanisms. Mice were fed on either a HP, HC, or HF diet for 5 weeks. (a) Proportion of B220^+^ B cells in the Peyer’s patches (n=8 per diet) and in the (b) mesenteric lymph nodes (n=4-7 per diet) were determined by flow cytometry. (c) Representative flow cytometry plot representing the proportion of B220^+^IgA^+^ plasmablasts in the Peyer’s patches and (d) corresponding scatter bar graph (n=8 per diet). (e) Representative flow cytometry plot of and graph representing the proportion of CD95^+^GL7^+^ germinal centre B cells in the Peyer’s patches and (f) corresponding scatter bar graph (n=8 per diet). Data are represented as mean ± SEM.

### High protein feeding modulates the lamina propria cytokine environment, specifically APRIL, to favour IgA production

T cell-independent induction of IgA CSR results from the complex interaction between gut microbes, host gut epithelial cells, immune cells and the stromal cells of the lamina propria. TLR activation in epithelial cells lead to the production of A Proliferation-inducing Ligand (APRIL) and B cell activating factor (BAFF), the major cytokines in B cell IgA CSR, as well as thymic stromal lymphopoietin (TSLP). TSLP further amplifies this signal by inducing APRIL and BAFF production by dendritic cells^13^.

To determine whether diet composition affects these cytokines, we quantified their expression levels in the small intestines of mice fed on HP, HC and HF diets by qPCR. The expression of *April* was significantly higher under HP feeding conditions, approximately 2-fold greater than HC and HF fed mice (Fig. 3a). Similarly, *Baff* was highly expressed under HP feeding conditions, compared to HF feeding conditions but HC-fed mice had levels of *Baff* expression similar to HP-fed mice (Fig. 3b). This suggests that increased expression of *April* might account for elevated sIgA levels under HP feeding conditions. Like *Baff, Tslp* expression was significantly lower in HF fed mice, whilst HP and HC fed mice had similar higher levels of expression (Fig. 3c). TSLP biases T cell differentiation towards Th2 T cells which are characterised by their production of the cytokine IL-4. IL-4 has been shown to promote IgA CSR as well as pIgR expression. Indeed, mice fed on a HP diet had significantly elevated expression of *Il4*, while HF fed mice had the lowest expression (Fig. 3d). Among other key cytokines involved in IgA CSR induction, IL-10 and TGF-beta produced by epithelial cells or dendritic cells are co-signals necessary for BAFF and APRIL to mediate their effects. Both *Il10* and *Tgfb* were expressed at highest levels in HP fed mice and consistently at lowest levels in HF fed mice (Fig. 3e-f).

**Figure 3:**
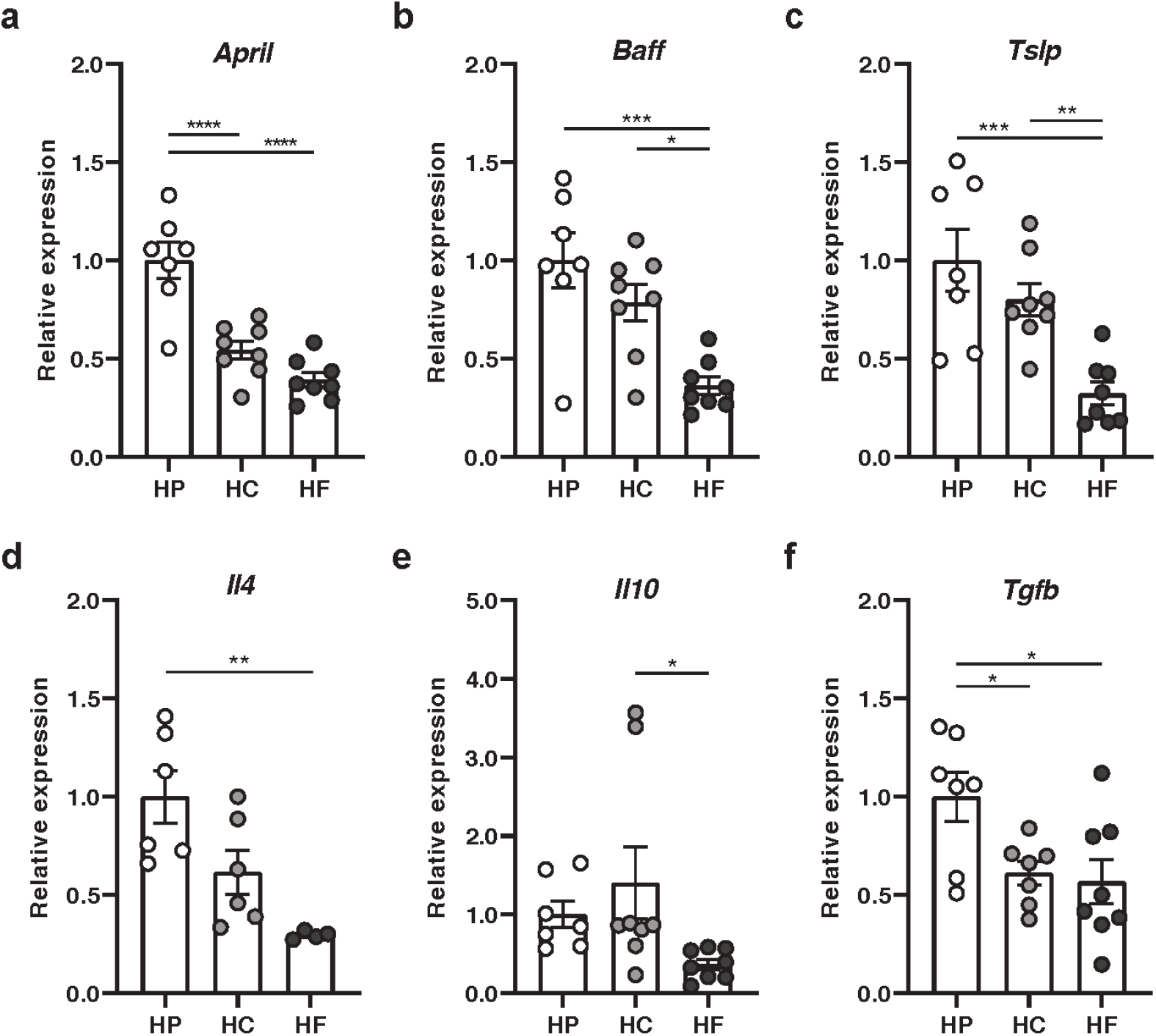
High protein feeding promotes a pro-IgA cytokine environment, specifically APRIL in the lamina propria. Mice were fed on either a HP, HC or HF diet for 5 weeks and intestinal ileal gene expression of (a) April, (b) Baff, (c) Tslp, (d) Il4, (e) Il10 and (f) Tgfb was determined by qPCR (n=6-8 per diet). Data are represented as mean ± SEM.

Together, we demonstrate that HP feeding induces the highest expression of cytokines involved in IgA CSR, and that these key cytokines are lowest under HF feeding.

### Dietary intervention significantly affects the small intestine microbiota composition

Diet is one of the most influential factors driving gut microbiota composition, which in turn modulates host IgA responses. To characterise the impact of HP, HC and HF diet feeding on the small intestinal gut microbiota composition, we performed 16S rRNA DNA sequencing of small intestine luminal samples. Analysis demonstrated equal sequencing reads between samples as well as adequate sequencing depth for all samples (Supplementary Table 5 and Supplementary Fig. 3a-b).

The diversity of the gut microbiota is often used as a marker of a “healthy microbiome”. We found that while bacterial richness was similar across all diets (Fig. 4a), HF feeding results in lower diversity measures such as evenness (Fig. 4b) and Inverse Simpson index (Fig. 4c), suggesting a dysbiotic small intestinal microbiome. Principal coordinates analysis (PCoA) of UniFrac distances revealed mice fed on each diet had significantly distinct gut microbiota compositions, with HF feeding having the most distinct bacterial communities in both unweighted UniFrac PCoA (Fig. 4d) and weighted UniFrac PCoA (Fig. 4e) analyses.

**Figure 4:**
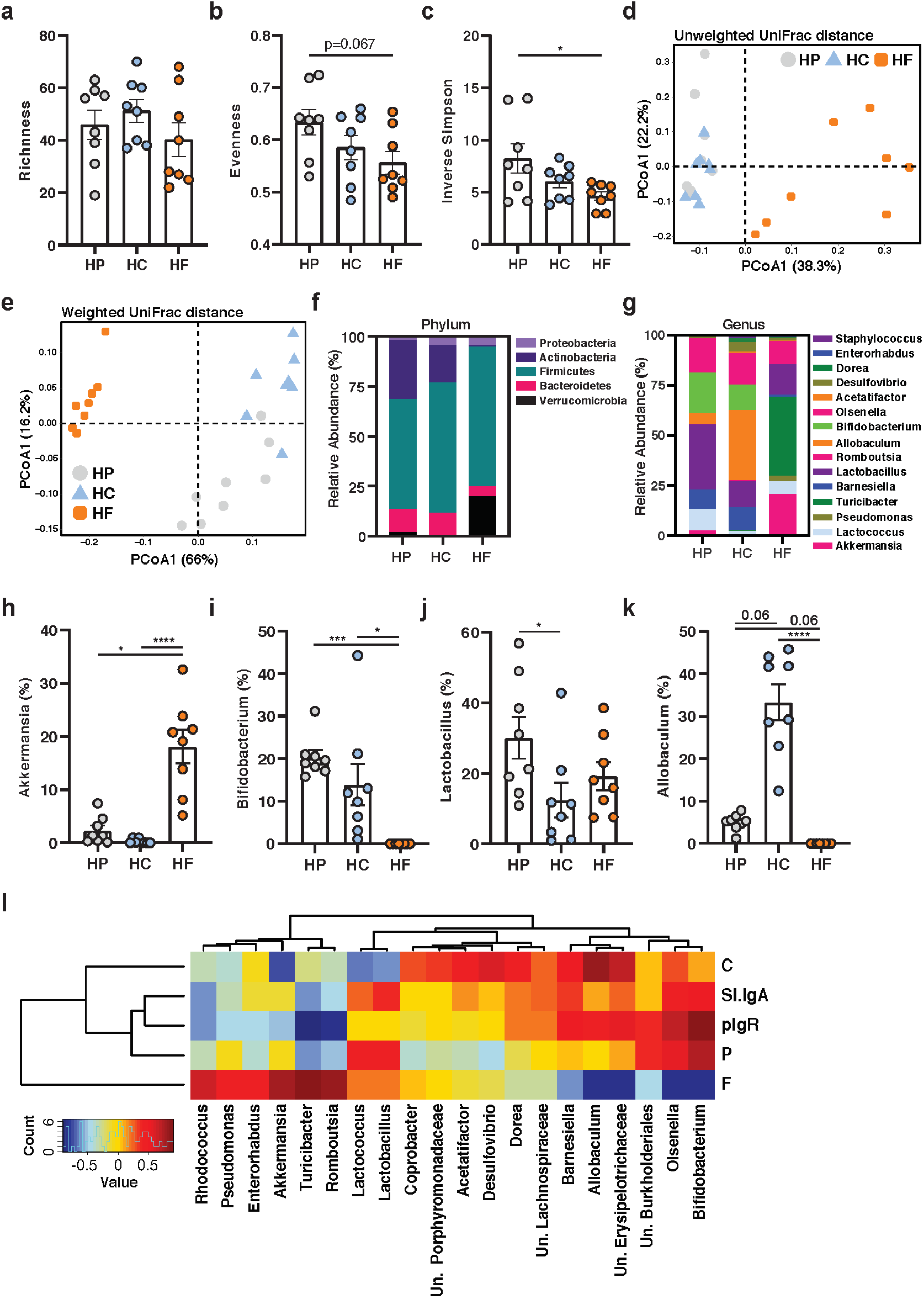
Dietary intervention significantly affects the small intestine microbiota composition. Mice were fed on either a HP, HC or HF diet for 5 weeks and DNA was extracted from small intestinal content for 16S rRNA sequencing (n=8 per diet). Diversity of the small intestinal microbiome was determined by (a) Richness, (b) Evenness and (c) Inverse Simpson Index. Differences in the structure of the small intestinal microbiota communities were determined by principal coordinate analysis of both (d) Unweighted and (e) Weighted UniFrac distances. Relative abundance of bacteria in the small intestine represented at the (f) phylum and (g) genus level (showing only the top 15 genera). Relative abundance of (h) Akkermansia, (i) Bifidobacterium, (j) Lactobacillus and (k) Allobaculum. (l) Spearman’s correlation representing the association between small intestinal microbiome at the genus level, small intestinal IgA levels and macronutrient composition in diet. Data are represented as mean ± SEM.

At the phylum level, HF feeding was associated with a microbiota with a higher ratio of *Firmicutes:Bacteroidetes* than the microbiota of HP- and HC-fed mice (Fig. 4f). The increase in *Firmicutes* is likely due to the overrepresentation of bacteria from the genus *Turicibacter* and *Staphylococcus*, while the abundance of *Bernesiella* from the phylum *Bacteroidetes* were underrepresented (Fig. 4g). The other main characteristic of a HF-fed microbiota is the overrepresentation of bacteria of the phylum *Verrucomicrobia*, particularly from the genus *Akkermansia* (Fig. 4h). While increased *Proteobacteria* have been linked to inflammation^14^, this phylum had the lowest abundance in HP-fed mice (Fig. 4f). The microbiota of the HP-fed mice was surprisingly characterised by the increased abundance of bacteria considered as probiotics with anti-inflammatory properties such as *Bifidobacterium* (Fig. 4i), *Lactobacillus* and *Lactococcus* (Fig. 4g, j). Finally, HC feeding was characterised by the over-representation of bacteria from the genus *Allobacullum* (Fig. 4g, k). By Spearman’s correlation, we confirmed that dietary protein content was more associated to sIgA than carbohydrate and fat (Fig. 4l). Interestingly, both protein and sIgA were highly associated with *Lactococcus* and *Lactobacillus* (which is under-represented in a HC diet), as well as *Olsenella* and *Bifidobacterium* (which is underrepresented in a HF diet). Overall, macronutrient composition significantly impacts both the alpha and beta diversity of the small intestine microbiome, with fat appearing to be the dominant driver.

### EV derived from high protein-fed microbiota activate epithelial TLR4 and promote the expression of PIGR and APRIL

While we identified that each diet had an impact on the gut microbiota, we then investigated the mechanisms through which the HP microbiota promote the host sIgA response. We first determined whether the intestinal environment of HP-fed mice could differentially affect TLR4 signalling. TLR4 is one of the main TLRs expressed by the small intestinal enterocytes and is involved in IgA CSR ^5^. HEK 293 cells transfected with TLR4 (HEK-Blue mTLR4) were incubated with small intestine contents isolated from mice fed on HP, HC or HF diets. HEK cells stimulated with HP small intestine content had significantly highest levels of TLR4 activation (Fig. 5a) suggesting the highest potential for HP fed mice microbiota to activate host cells.

**Figure 5:**
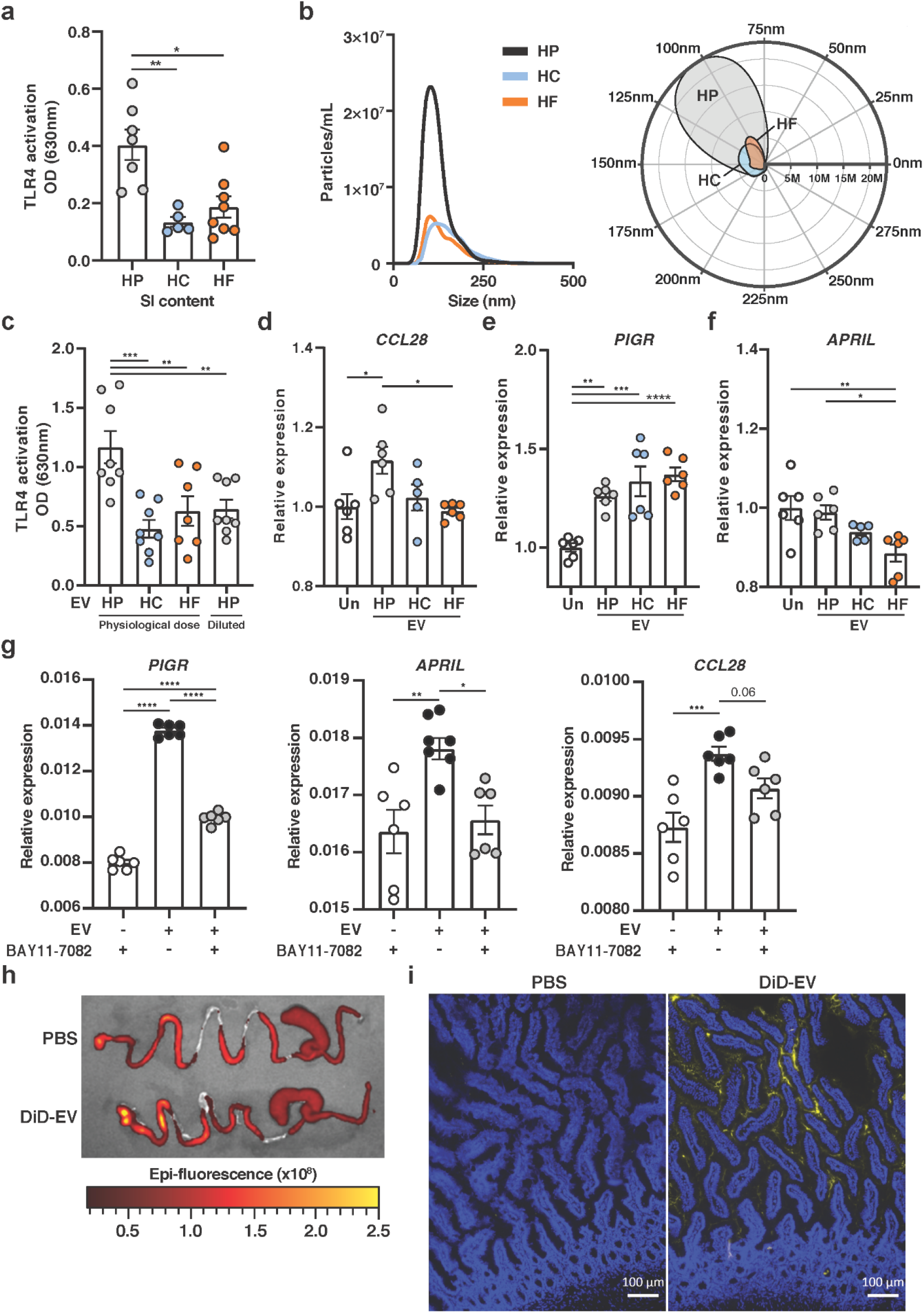
EV derived from high protein-fed microbiota activate epithelial TLR4 and promote the expression of PIGR and APRIL. Mice were fed on either a HP, HC, or HF diet for 5 weeks. (a) Small intestinal content (1:100 dilution) (n=5-8 per diet) was incubated overnight with HEK-TLR4 cell line and TLR4 activation was quantified at 630nm. (b) Cecum microbiota-derived EV were characterised by NTA (n=2 per diet pooled from n=4 mice each) and represented as an XY plot (left, 0-500nm) or polar plot (right, angular axis represents particle size between 0-300nm and radial axis represents particle concentration in number/mL). (c) Microbiota-derived EV were incubated at physiological ratio (approximately 3:1:1 of HP:HC:HF), or at 1:1:1 ratio (HP diluted) overnight with HEK-TLR4 cell line and TLR activation measured at 630nm (n=7-8 per condition). (D-F) HT-29 were stimulated with microbiota-derived EV for 16h and expression of (d) CCL28, (e) PIGR and (f) APRIL were quantified by qPCR (n=6 per group) as well as (g) in the presence or absence of 10µM of NF-κB inhibitor BAY 11-7082, or BAY 11-7082 alone (n=6 per condition). (h) Representative ex vivo epi-fluorescence imaging of gastrointestinal tract 6 hours post-oral administration of DiD-stained EV or PBS. (i) Presence of DiD-EV (yellow) and DAPI (blue) were assessed by fluorescent microscopy from sections of small intestine isolated from mice 6 hours after oral administration, with PBS as control. Scale bar represents 100µm. Data are represented as mean ± SEM.

Typically, the mucus layer mediates an efficient physical separation of microbes from the gut epithelium. We therefore hypothesised that *in vivo* activation of TLR4 might be mediated by smaller structures containing PAMPs, rather than through physical interaction with whole bacteria. Bacterial EV are small spherical structures below 300nm diameter in size, produced by the budding of the membrane and consequently contain PAMPs. EV can cargo nucleic acid, protein, metabolites and other molecules and are key component of bacterial communication^15^. EV derived from *Bacteroides fragilis* have been shown to promote regulatory T cell differentiation through the stimulation of TLR^16^, highlighting the ability of EV to traverse the mucus layer to reach the host. As such, we investigated whether EV derived from the microbiota of HP-fed mice might activate TLR and promote cytokines involved in IgA CSR.

We first characterised the number and size distribution of microbiota-derived EV from mice fed on the different diets by nanoparticle tracking analysis (NTA). The size distribution of the microbiota-derived EV was within the expected range of 50-300nm (Fig. 5b). While the size distribution was similar across the diets, there was approximately a 3-fold increase in the concentration of microbiota-derived EV under HP feeding conditions (Fig. 5b).

To determine whether HP-derived microbiota differentially activate TLR4 via its EV, we incubated TLR4 transfected HEK cells with purified EV isolated from microbiota of mice fed on the different diets. We first incubated the HEK-TLR4 cells with relative doses of EV similar to those observed *in vivo*. Consistent with the previous results showing highest potential of small intestine content to activate TLR4, we found that EV derived from HP microbiota stimulated TLR4 and to the highest extent (Fig. 5c). However, when HEK cells were incubated with the same concentration of EV from each group, the activation of TLR4 was similar. Thus, the higher potential of HP EV to activate TLR4 was due to a quantitative rather than a qualitative effect. To determine their potential impact on host sIgA production and translocation, we quantified the gene expression of *CCL28, APRIL* and *PIGR* in HT29 cells incubated with the EV. EV derived from HP microbiota significantly upregulated *CCL28*, compared to unstimulated cells or to HF EV (Fig. 5d) and we observed the same trend when compared to HC EV. EV from all groups upregulated the expression of *PIGR* (Fig. 5e). Interestingly, while HP did not affect *APRIL*, HF EV significantly decreased its expression with HC EV showing a similar trend (Fig. 5f). To confirm whether EV mediated these effects via TLR signalling, we cultured HT-29 with EV in the presence of BAY 11-7082, an inhibitor of NF-κB involved in TLR signalling. Addition of BAY 11-7082 abrogated the effect of EV on the expression of *PIGR, APRIL* and *CCL28* (Fig. 5g) suggesting that EV mediated their effects via TLR signalling.

To determine whether microbiota-derived EV could reach the host small intestine epithelium *in vivo*, we isolated microbiota-EV from mice fed on normal chow, fluorescently labelled them with DiD, and administered them by gavage to another set of mice. *Ex vivo* imaging of gut tissue by IVIS revealed higher epi-fluorescence intensity in the gastrointestinal tract of mice receiving DiD-EV compared to PBS control showing that DiD-EV readily reached the small intestine (Fig. 5h). There was presence of DiD-EV in both the lumen and mucosa of these mice, as determined by fluorescence microscopy (Fig. 5i).

Together, these data highlight a close interaction between bacterial EV and host cells *in vivo* as well as a novel role for gut microbiota-derived EV on the regulation of host genes involved in sIgA regulation via TLR activation.

### Bacterial metabolites do not affect IgA-related genes but promote bacterial ROS and bacterial EV production

In addition to TLR activation, the gut microbiota can also modulate host sIgA production through the generation of metabolites, such as short-chain fatty acids^17–19^. To determine whether such mechanisms are involved, we quantified microbiota-derived metabolites by NMR and focused on metabolites that are significantly upregulated under HP feeding conditions. We found that succinate (Fig. 6a), pyruvate (Fig. 6b) and propionate (Fig. 6c) were significantly elevated in HP, compared to mice fed on HC and HF diets. To determine their effects on *CCL28, APRIL and PIGR* expression, we incubated HT-29 cells with each of these candidate metabolites. We found that, unlike EV-stimulated HT-29, neither pyruvate, propionate nor succinate could directly induce the expression of *CCL28, APRIL or PIGR* expression (Fig. 6d-f). As such, we hypothesized that changes to the gut metabolite environment would affect the gut microbiota, rather than the host directly to elicit host sIgA response.

**Figure 6:**
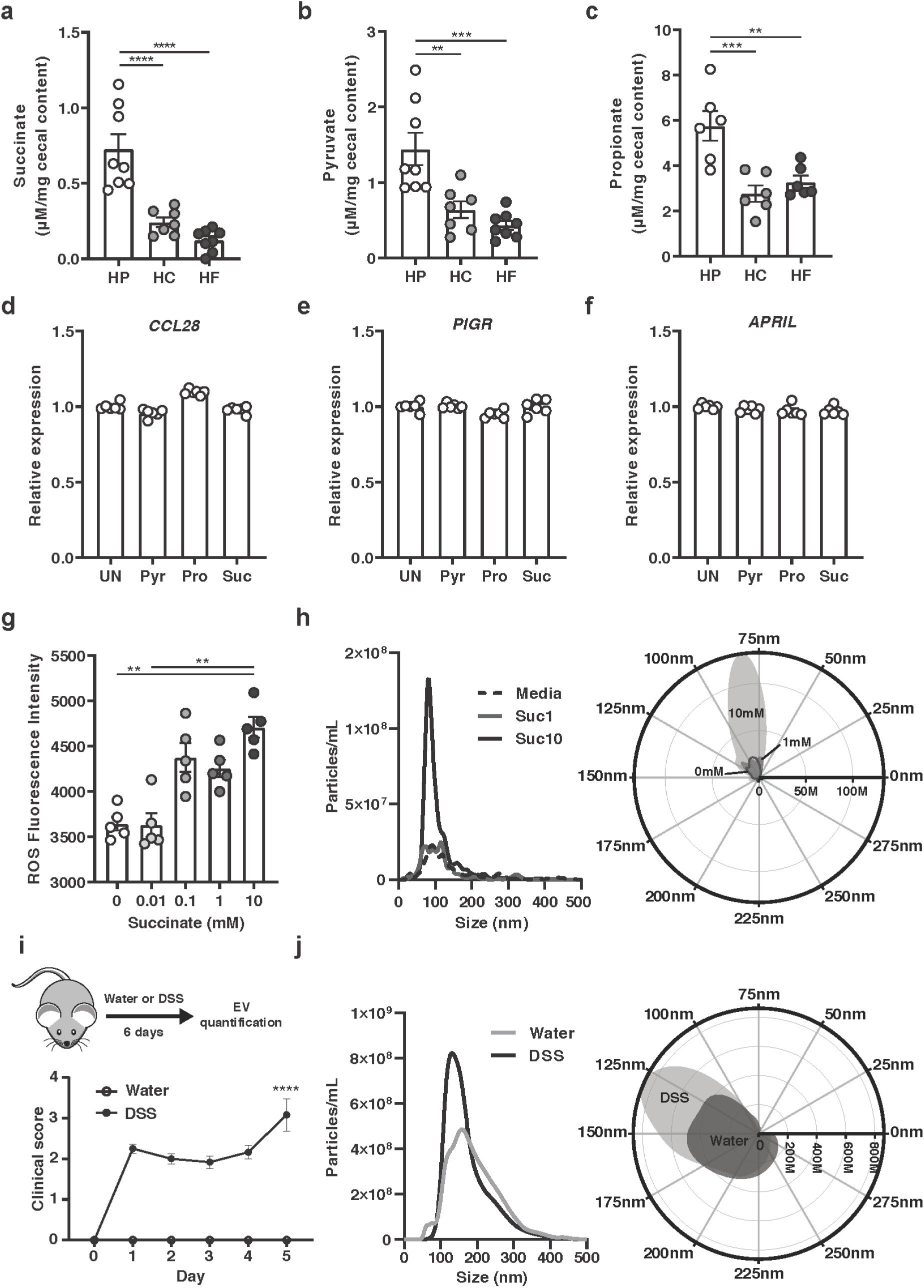
Bacterial metabolites do not affect IgA-related genes but promote bacterial ROS and bacterial EV production. Metabolites from cecal content of mice fed on a HP, HC or HF diet for 5 weeks was quantified by NMR spectroscopy for (a) succinate, (b) pyruvate and (c) propionate (n=7-8 per diet). (d-f) HT29 were stimulated overnight with 1mM of pyruvate (Pyr), propionate (Pro) or succinate (Suc) and gene expression levels of (d) CCL28, (e) PIGR and (f) APRIL quantified by qPCR (n=6 per condition). (g) E. coli was grown in the presence of increasing concentration of succinate (0-10mM) for 2h and ROS production quantified by the conversion of 2’,7’-dichlorofluorescein diacetate to 2’,7’-dichlorofluorescein (n=5 per condition). (h) E. coli was grown in the presence of succinate (0, 1 or 10mM) for 16h and EV isolated and quantified by NTA (n=5-6 per condition) and represented as scatter bar plot (left) or polar plot (right, angular axis represents particle size between 0-300nm and radial axis represents particle concentration in number/mL). (i) Mice received 3% DSS in drinking water for 6 days (upper) and colitis development was scored daily (lower) (n=6 per group). (i) At endpoint, faecal EV were isolated and quantified by NTA (n=6 per condition) and represented as an XY plot (left, 0-500nm) or polar plot (right, angular axis represents particle size between 0-300nm and radial axis represents particle concentration in number/mL). Data are represented as mean ± SEM.

Like any cell types, bacteria can produce EV in response to stress signals such as reactive oxygen species (ROS)^15^. Succinate, which we found to be highly upregulated in HP-fed mice, has been shown to increase ROS in host cells such as macrophages^20^. In contrast, pyruvate and propionate were shown to have anti-oxidative effects^21,22^. High levels of microbiota-produced succinate under HP-feeding conditions might promote bacterial ROS, which would in turn stimulate the release of EV. To test this hypothesis, we incubated *E. coli* with increasing concentrations of succinate and quantified ROS. Strikingly, succinate increased ROS in a dosedependent manner (Fig. 6g) without affecting bacterial growth (Supplementary Fig. 4a). Conversely, neither propionate nor pyruvate increased ROS levels (Supplementary Fig. 4b). To determine the impact of succinate on EV production, we cultured *E. coli* with succinate for 16h and found that succinate significantly increased EV production by NTA (Fig. 6h) while propionate and pyruvate did not (Supplementary Fig. 4c). These results uncover a novel mechanism through which gut bacterial vesiculation is regulated by the metabolites in the environment, particularly succinate. Increased succinate has also been reported in human inflammatory bowel disease as well as in DSS-induced colitis in mice^23^. In both cases, increased TLR activation by microbial PAMPs has been reported in the gut^24^. Based on our results, we hypothesize that high succinate in colitis will correlate with higher gut bacterial EV production that may further potentiate gut inflammation by increasing TLR activation. To test whether DSS colitis could affect bacterial EV production, we treated mice for 6 days with 3% DSS in drinking water and quantified the production of faecal EV by NTA (Fig. 6i). As expected, mice treated on DSS had higher clinical score (Fig. 6i). While the size distribution of EV isolated from mice with colitis was similar to those isolated from control mice, the quantity of EV was almost twice higher in the DSS mice (Fig. 6j). Altogether, these results highlight a novel role for bacterial metabolite succinate in the induction of bacterial EV that can in turn activate TLR signalling and modulate host gene expression.

## Discussion

In the present study, we have identified a key role for dietary macronutrient composition on the dynamic regulation of IgA production and sIgA release in the small intestinal lumen. Dietary protein was the major driver of T cell-independent sIgA response, associated with increased expression of the CSR promoting cytokines APRIL and BAFF. Increased luminal sIgA was associated with elevated expression of the transporter *pIgR* that was correlated with the highest expression of IL-4, a cytokine involved in both IgA induction and pIgR expression. These changes are commonly observed under pro-inflammatory conditions linked to increased activation of host pattern-recognition receptors by gut bacterial products. Accordingly, we found that intestinal luminal content from HP-fed mice activated TLR4-NFκB pathway to the highest extent compared to contents obtained from HC and HF-fed mice. This was attributed to the greater production of microbiota-derived EV under HP feeding conditions, that could activate TLR4 signalling pathway. Increased production of EV is likely the result of increased microbial stress, as previously reported in bacteria^15^. Accordingly, we found that high protein consumption significantly increased microbial succinate production, which we demonstrated to promote bacterial ROS production in a concentration-dependent manner *in vitro*. This work highlights a novel mechanism through which the gut microbiota regulates host sIgA response modulated by diet composition.

The majority of sIgA produced under homeostatic conditions is induced by commensal bacteria via T cell-independent mechanisms. This appears to be the default pathway as sIgA induced by the monocolonization of germ-free mice with various strains of bacteria were mostly non-specific and polyreactive in nature^25^. This observation highlights that the presence, rather than the composition of microbiota is important in inducing sIgA and the mechanism likely involves a shared feature of bacteria, such as their capacity to activate host TLR via MAMPs. Accordingly, our small intestinal microbiome data revealed no obvious dysbiosis associated with HP feeding that could explain the increase of sIgA. Rather, a HP microbiome was associated with a higher abundance of bacteria from the genus *Lactobacillus* and *Bifidobacterium*, commonly considered as probiotics with purported health benefits. Despite this, a HP microbiome had an exacerbated pro-inflammatory potential via increased production of EV that can activate TLR4. How the gut microbiota can adequately activate TLR under homeostatic conditions remains unclear, as TLR is usually inaccessible to whole bacteria due to the presence of a mucus layer. We highlight that microbiota-derived EV might be the link as they can traverse the mucus and are able to activate TLR4 as well as IgA-related epithelial genes *in vitro*, revealing a new mechanism underpinning host microbiota-mutualism. HT-29, being a human cell line, highlights the translational potential of our finding. Quantification of microbiota-derived EV could prove a useful tool to evaluate the functional characteristics as well as the inflammatory potential of the gut microbiota as an alternative to sequencing.

Our study examined both qualitative and quantitative effects of microbiota-derived EV on the host. TLR4 and CCL28 were modulated by EV in a dose-dependent manner, with HP feeding conditions increasing the activity of the former and gene expression of the latter. This suggests that EV could be a mechanism by which the host perceives changes in the state of the gut microbiota via TLR activation. As a result, the host would mobilize B cells to the gut and increase the translocation of sIgA to control the situation based on the amount of EV sensed. This could have clinical implications in diseases such as food allergy, in which low levels of IgA are commonly reported. We also uncovered a qualitative effect of EV on the host. Significant changes in the gut microbiota composition have been shown to promote the production of different types of EV. A complete analysis of the features of EV derived from HP, HC and HF microbiota, outside the scope of this study, would confirm this. Expression of *April* was not upregulated by high EV levels under HP feeding but was rather inhibited by HF EV suggesting a specific regulation of host genes by bacterial EV. This result aligned with the very low levels of sIgA observed in the gut of HF fed mice and suggests that EV can regulate the host through both TLR-dependent and TLR-independent mechanisms.

The high levels of succinate under HP-feeding conditions remain unexplained. We could not identify a specific microbiome signature that would account for greater succinate production in HP-fed mice. One possibility is the disappearance of succinate consumers in the HP microbiota. Our finding that succinate could modulate ROS production in bacteria is novel and the long-term impact of this on the gut microbiota yet to be determined. ROS is known to induce bacterial mutation and antibiotic resistance which could thus raise concerns that high protein diet feeding could promote the potential growth of antibiotic-resistant bacteria. Moreover, we identified higher levels of bacterial EV in DSS colitis, a disease in which gut bacterial dysbiosis, as well as high succinate levels, were previously reported^23^. This result suggests that dysbiosis, in general, could be sensed by the host via EV yet whether bacterial dysbiosis in other context than IBD is associated with higher EV production remains unknown.

In summary, we describe a new pathway by which the gut microbiota can regulate T cell-independent IgA induction via bacteria-derived EV. More broadly, we show a novel mechanism through which the microbiota impact the host. This opens exciting new prospects of characterizing microbiota EV as a biomarker for dysbiosis, as well as the potential use of microbiota EV as a postbiotic to restore gut homeostasis.

## Methods

### Animals and housing

Male C57BL/6 mice (6 weeks of age) were purchased from Animal Bioresource (NSW, Australia) and maintained under specific pathogen-free conditions in the animal facility of the Charles Perkins Centre. All experiments were performed in accordance to protocols approved by the University of Sydney Animal Ethics Committee (Protocol 2017/1280 and 2019/1493).

### Animal diets and feeding

Animals were fed ad libitum on either a high protein, moderately low carbohydrate, moderately low-fat diet (P60, C20, F20; henceforth, HP); a low protein, high carbohydrate, moderately low-fat diet (P5, C75, F20; henceforth, HC), or a low protein, moderately low carbohydrate, high-fat diet (P5, C20, F75; henceforth, HF). These three diets represented the extremes (apices) from a larger set of 10 diet compositions (Supplementary Table 1) that were used in a subset of the experiments, encompassing a macronutrient range of protein (5-60%), carbohydrate (20-75%), and fat (20-75%) chosen using nutritional geometry to comprehensively sample dietary macronutrient mixture space^9^. All diets were isocaloric at 14.5 MJ/kg and based on modification to the semi-purified AIN93G formulation and purchased from Specialty Feeds, Glenn Forest, Australia (SF17-188-SF17-197).

### Quantification of immunoglobulins

Small intestinal content was resuspended in PBS (100mg/ml) containing cOmplete™ Protease Inhibitor Cocktail (Roche), centrifuged at 14,000xg for 10min to pellet bacteria and debris and supernatant collected. Quantification of IgA and IgM level was performed using the Mouse IgA or IgM ELISA quantification set (Bethyl Laboratories).

### Immunofluorescence

For immunohistochemistry analysis, frozen small intestine sections were fixed with ice-cold methanol, and incubated with goat anti-mouse IgA (Bethyl Laboratories) followed by AF488-conjugated chicken anti-goat IgG secondary antibody (Invitrogen). Sections were stained with DAPI and examined with an Olympus BX51 Microscope and images captured using the cellSens Standard software.

### Flow cytometry

Single-cell suspension were prepared by mechanical disruption of tissues through a 100µm filter. Cells were incubated with LIVE/DEAD™ Fixable Blue Dead Cell Stain Kit (ThermoFisher Scientific) for exclusion of non-viable cells and anti-mouse CD16/32 (91; BioLegend) for blocking non-specific binding, then stained with the antibodies described in Supplementary Table 6 and analysed on the LSRII (BD Bioscience). Data were analysed with the FlowJo v10.4.2 software (FlowJo LLC).

### RNA extraction and qPCR

Total RNA was extracted using TRI Reagent (Sigma Aldrich) following the manufacturer’s instructions and cDNA generated using the High-Capacity cDNA Reverse Transcription Kit (ThermoFisher Scientific). qPCR was conducted using the Power SYBR™ Green PCR Master Mix (ThermoFisher Scientific) on a LightCycler® 480 Instrument II (Roche Life Science). Primer sequences are provided in Supplementary Table 7.

### 16S rRNA gene sequencing and bioinformatics

DNA from small intestinal jejunum content was extracted using the FastDNA™ SPIN Kit for Feces (MP Biomedicals). Illumina sequencing of the V4 region (515f-806r) of the 16S rRNA gene was performed commercially at the Ramaciotti Centre for Genomics (UNSW, Australia). Paired-end reads (2×250bp) were processed with the dada2 package (1.12.1) using R software (3.6.1) and taxonomy was assigned using the Ribosomal Database Project training set with specie level taxonomy assignment (doi:10.5281/zenodo.801828). Further analyses were performed with the R packages phyloseq (1.28.0) and vegan (2.5.5). Gene sequencing data were deposited in the European Nucleotide Archive under accession number PRJEB39583.

### EV isolation and nanoparticle tracking analysis (NTA)

100mg of cecum content was homogenized in 0.02µm filtered PBS and centrifuged at 340xg for 10min at 4°C. Supernatant was then centrifuged at 10,000xg for 20min at 4°C followed by 18,000xg for 45min at 4°C, filtered through a 0.22µm filter and centrifuged at 100,000xg for 2hr at 4°C. EV pellet was resuspended in 0.02µm filtered PBS. EV particle size and concentration were assessed by NTA on a Nanosight NS300 (Malvern Instruments Limited). NTA 3.2 software was used to calculate EV concentration and size distribution.

### Ex vivo imaging of EV uptake

Fluorescent staining of EV was performed with the Vybrant DiD Cell Labelling Kit (Invitrogen) at a final concentration of 5µM for 30min at room temperature and washed with PBS by ultracentrifugation at 100,000xg for 2h at 4°C. Animals were intragastrically administered with 80µg of DiD-EV and organs imaged 6h later with the IVIS Spectrum In Vivo Imaging System (Perkin Elmer) at excitation:640nm and emission:680nm. Image was analysed using the Living Image 4.5 Software (Perkin Elmer) and fluorescence was quantified as radiant efficiency.

### TLR4 activity assay and cell culture

HEK-Blue mTLR4 reporter cell line (InvivoGen) was maintained in complete DMEM (Gibco) containing 10% FBS (Bovogen Biologicals), 100U/ml Penicillin and 100µg/ml Streptomycin (Gibco), 2mM L-glutamine (Sigma-Aldrich) with 100µg/ml Normocin and 1X HEK-Blue selection (InvivoGen). TLR4 activation by bacterial-derived EV or diluted small intestinal content was assessed according to the manufacturer’s instructions. HT-29 cells (ATCC) were maintained in complete DMEM media and stimulated with small intestinal content or bacteria-derived EV for 16h. NF-κB inhibitor BAY 11-7082 (Sigma-Aldrich) was used at 10µM.

### Detection of polar metabolites

For detection of polar metabolites by ^1^H nuclear magnetic resonance spectroscopy, cecal content was homogenized in deuterium oxide (Sigma-Aldrich) at a concentration of 100mg/ml and centrifuged at 14,000xg for 5min at 4°C. Resulting supernatant was diluted with sodium triphosphate buffer (pH 7.0) containing 0.5mM 4,4-dimethyl-4-silapentane-1-sulfonic acid as internal standard (All from Sigma-Aldrich). Samples were run on a Bruker 600 MHz AVANCE III spectrometer and resulting spectra processed and analyzed using the Chenomx NMR Suite v8.4 software (Chenomx Inc).

### Bacterial culture and ROS quantification

*Escherichia coli* (K-12 MG1655) were cultured in Luria-Bertani broth with succinate, propionate or pyruvate at indicated doses and supernatants collected for EV quantification 16h later. For quantification of ROS production, *E. coli* were treated with succinate at indicated doses for 2h, and cells pelleted by centrifugation (5000rpm for 5min). Pellet was resuspended in PBS containing 10µM of 2’,7’-dichlorofluorescein diacetate (Sigma-Aldrich) and incubated for 60min at room temperature. Fluorescence intensity was measured using a TECAN Infinite M1000 microplate reader (Excitation 485nm, Emission 538nm, Gain 100).

### Experimental model of colitis

Murine model of colitis was induced by the administration of 3% dextran sodium sulfate (DSS, MW: 36,000-50,000, MP Biomedicals) in drinking water for 6 days. Mice were monitored daily and clinical score was assessed as follows: 0-normal faeces; 1-soft faeces; 2-pasty faeces; 3-liquid faeces; 4-moderate rectal bleeding; 5-severe rectal bleeding; 6-haemorrhagia/diarrhoea and 7-signs of mobidity.

### Mixture model and statistical analyses

The effects of dietary macronutrient composition on outcomes were analysed using mixture models (also known as Scheffé’s polynomials). Analyses were performed on each outcome variable separately. Models were implemented using the *mixexp* package (1.2.5) using R software (3.6.1). For each outcome, four models equivalent to equations described by Lawson and Willden^10^ as well as a null model was fitted. Collectively, these models test for no effect, linear effects, and non-linear effects of macronutrient composition on the outcome variable. The most appropriate model was determined by Akaike information criterion (AIC), where the lowest AIC value (where AIC was within 2 points of difference, the simpler model was selected) was deemed the most appropriate model. Predictions from the best model was represented on a right-angled mixture triangle plot^11^. For comparison between groups, Ordinary one-way ANOVA followed by Tukey’s multiple comparisons test was used. The differences were considered as significant when p<0.05 where *p<0.05, **p<0.01, ***p<0.001 and ****p<0.0001.

## Acknowledgments

This project was funded by the Australian Research Council grant APP160100627 and by the Sydney Medical School MCR BioMed-Connect Grants. We thank Alexandra Angelatos for her help with some aspect of the project.

## Author Contributions

J.T performed most of the experiments, participated to the project design and wrote the manuscript, D.N, J.T and G.V.P participated to the experiments, A.S and M.R. helped with the data analysis, J.W and N.J.C.K., G.E.G. and S.J.S. helped with the study design, L.M participated to the study design, supervised the study and wrote the manuscript.

## Data availability

The data supporting the findings of this study are available from the corresponding author upon reasonable request

## References

1. Sommer, F. & Bäckhed, F. The gut microbiota--masters of host development and physiology. Nat. Rev. Microbiol. 11, 227–238 (2013).

2. Yel, L. Selective IgA Deficiency. J Clin Immunol 30, 10–16 (2010).

3. Fagarasan, S., Kawamoto, S., Kanagawa, O. & Suzuki, K. Adaptive immune regulation in the gut: T cell-dependent and T cell-independent IgA synthesis. Annu. Rev. Immunol. 28, 243–273 (2010).

4. Macpherson, A. J. et al. A primitive T cell-independent mechanism of intestinal mucosal IgA responses to commensal bacteria. Science 288, 2222–2226 (2000).

5. Shang, L. et al. Toll-like receptor signaling in small intestinal epithelium promotes B-cell recruitment and IgA production in lamina propria. Gastroenterology 135, 529–538 (2008).

6. He, B. et al. Intestinal bacteria trigger T cell-independent immunoglobulin A(2) class switching by inducing epithelial-cell secretion of the cytokine APRIL. Immunity 26, 812–826 (2007).

7. Hapfelmeier, S. et al. Reversible microbial colonization of germ-free mice reveals the dynamics of IgA immune responses. Science 328, 1705–1709 (2010).

8. Lubomski, M. et al. Parkinson’s disease and the gastrointestinal microbiome. J. Neurol. (2019) doi:10.1007/s00415-019-09320-1.

9. Simpson, S. J., Le Couteur, D. G. & Raubenheimer, D. Putting the balance back in diet. Cell 161, 18–23 (2015).

10. Lawson, J. & Willden, C. Mixture Experiments in R Using mixexp. Journal of Statistical Software 72, 1–20 (2016).

11. Raubenheimer, D. Toward a quantitative nutritional ecology: the right-angled mixture triangle. Ecological Monographs 81, 407–427 (2011).

12. Saner, C. et al. Evidence for Protein Leverage in Children and Adolescents with Obesity. Obesity (Silver Spring) 28, 822–829 (2020).

13. Wells, J. M., Rossi, O., Meijerink, M. & van Baarlen, P. Epithelial crosstalk at the microbiota-mucosal interface. Proc. Natl. Acad. Sci. U.S.A. 108 Suppl 1, 4607–4614 (2011).

14. Shin, N.-R., Whon, T. W. & Bae, J.-W. Proteobacteria: microbial signature of dysbiosis in gut microbiota. Trends Biotechnol. 33, 496–503 (2015).

15. Macia, L., Nanan, R., Hosseini-Beheshti, E. & Grau, G. E. Host- and Microbiota-Derived Extracellular Vesicles, Immune Function, and Disease Development. Int J Mol Sci 21, (2019).

16. Shen, Y. et al. Outer membrane vesicles of a human commensal mediate immune regulation and disease protection. Cell Host Microbe 12, 509–520 (2012).

17. Kim, M., Qie, Y., Park, J. & Kim, C. H. Gut Microbial Metabolites Fuel Host Antibody Responses. Cell Host Microbe 20, 202–214 (2016).

18. Tan, J. et al. Dietary Fiber and Bacterial SCFA Enhance Oral Tolerance and Protect against Food Allergy through Diverse Cellular Pathways. Cell Rep 15, 2809–2824 (2016).

19. Sanchez, H. N. et al. B cell-intrinsic epigenetic modulation of antibody responses by dietary fiber-derived short-chain fatty acids. Nat Commun 11, 60 (2020).

20. Mills, E. L. et al. Succinate Dehydrogenase Supports Metabolic Repurposing of Mitochondria to Drive Inflammatory Macrophages. Cell 167, 457-470.e13 (2016).

21. Tauffenberger, A., Fiumelli, H., Almustafa, S. & Magistretti, P. J. Lactate and pyruvate promote oxidative stress resistance through hormetic ROS signaling. Cell Death & Disease 10, 1–16 (2019).

22. Hoyles, L. et al. Microbiome-host systems interactions: protective effects of propionate upon the blood-brain barrier. Microbiome 6, 55 (2018).

23. Connors, J., Dawe, N. & Van Limbergen, J. The Role of Succinate in the Regulation of Intestinal Inflammation. Nutrients 11, (2018).

24. Lu, Y., Li, X., Liu, S., Zhang, Y. & Zhang, D. Toll-like Receptors and Inflammatory Bowel Disease. Front Immunol 9, 72 (2018).

25. Pabst, O. & Slack, E. IgA and the intestinal microbiota: the importance of being specific. Mucosal Immunol 13, 12–21 (2020).

