## Supplemental Information for "Dietary protein increases T cell independent sIgA production through changes in gut microbiota-derived extracellular vesicles"

**Supplementary Table 1 – Detailed composition of the experimental diets used in the study**

Diet formulation provided as gram of ingredients per 100g for each of the 10 diets used in the study. The diets are isocaloric at 14.5 MJ/kg with the following net metabolizable energy (NME) assigned: casein (13.3 kJ/g; 3.2 kCal/g), L-methionine (18 kJ/g; 4.3 kCal/g), Canola oil (36.6 kJ/g; 8.7 kCal/g), Wheat starch (14.13 kJ/g; 3.4 kCal/g), Dextrinized Starch (14.6 kJ/g; 3.49 kCal/g), Sucrose 14.92 kCal/g and cellulose was given a NME of 0 kCal/g.

| | | **Diet 1** | **Diet 2** | **Diet 3** | **Diet 4** | **Diet 5** | **Diet 6** | **Diet 7** | **Diet 8** | **Diet 9** | **Diet 10** |
|---|---|---|---|---|---|---|---|---|---|---|---|
| | **%P** | 60 | 5 | 5 | 33 | 33 | 5 | 14 | 14 | 42 | 24 |
| | **%C** | 20 | 75 | 20 | 47 | 20 | 47 | 29 | 57 | 29 | 38 |
| | **%F** | 20 | 20 | 75 | 20 | 47 | 48 | 57 | 29 | 29 | 38 |
| **INGREDIENTS** | | | | | | | | | | | |
| **Protein** | **Casein** | 64.87 | 5.04 | 5.04 | 35.50 | 35.50 | 5.04 | 14.83 | 14.83 | 45.29 | 25.71 |
| | **L-Methionine** | 0.30 | 0.30 | 0.30 | 0.30 | 0.30 | 0.30 | 0.30 | 0.30 | 0.30 | 0.30 |
| **Fat** | **Canola Oil** | 7.91 | 7.91 | 29.65 | 7.91 | 18.58 | 18.98 | 22.53 | 11.46 | 11.46 | 15.03 |
| **Carbohydrate** | **Wheat Starch** | 12.59 | 47.87 | 12.60 | 29.91 | 12.60 | 29.91 | 18.37 | 36.34 | 18.37 | 24.13 |
| | **Dextrinized Starch** | 4.10 | 15.57 | 4.10 | 9.73 | 4.10 | 9.73 | 5.98 | 11.82 | 5.98 | 7.85 |
| | **Sucrose** | 3.11 | 11.81 | 3.11 | 7.38 | 3.11 | 7.38 | 4.54 | 8.97 | 4.54 | 5.96 |
| **Minerals** | **$CaCO_3$** | 1.31 | 1.31 | 1.31 | 1.31 | 1.31 | 1.31 | 1.31 | 1.31 | 1.31 | 1.31 |
| | **NaCl** | 0.26 | 0.26 | 0.26 | 0.26 | 0.26 | 0.26 | 0.26 | 0.26 | 0.26 | 0.26 |
| | **AIN93 Trace Minerals** | 0.14 | 0.14 | 0.14 | 0.14 | 0.14 | 0.14 | 0.14 | 0.14 | 0.14 | 0.14 |
| | **$KH_2PO_4$** | 0.69 | 0.69 | 0.69 | 0.69 | 0.69 | 0.69 | 0.69 | 0.69 | 0.69 | 0.69 |
| | **$K_2SO_4$** | 0.16 | 0.16 | 0.16 | 0.16 | 0.16 | 0.16 | 0.16 | 0.16 | 0.16 | 0.16 |
| | **KCl** | 0.25 | 0.25 | 0.25 | 0.25 | 0.25 | 0.25 | 0.25 | 0.25 | 0.25 | 0.25 |
| | **$C_5H_{14}ClNO$** | 0.25 | 0.25 | 0.25 | 0.25 | 0.25 | 0.25 | 0.25 | 0.25 | 0.25 | 0.25 |
| **Vitamins** | **AIN93 vitamins** | 1.00 | 1.00 | 1.00 | 1.00 | 1.00 | 1.00 | 1.00 | 1.00 | 1.00 | 1.00 |
| **Cellulose** | **Cellulose** | 3.06 | 7.44 | 41.15 | 5.21 | 21.75 | 24.60 | 29.39 | 12.23 | 10.00 | 16.97 |

**Supplementary Table 2 – Statistical output for mixture models relating to sIgA**

| Model (Scheffé Polynomials) | Akaike Information Criterion | Degrees of freedom |
|---|---|---|
| 1 | 1168.363 | 4 |
| 2 | 1168.552 | 7 |
| 3 | 1156.861 | 11 |
| 4 | 1170.262 | 8 |
| Null | 1181.324 | 2 |

| **Model 1 Coefficients** | | | |
|---|---|---|---|
| **Components** | **Estimate (Std. Error)** | **t value** | **P(>\|t\|)** |
| Protein | 2094.2 (264.7) | 7.913 | 2.05e−11 |
| Fat | 517.6 (214.5) | 2.413 | 0.018334 |
| Carbohydrate | 868.8 (220.6) | 3.939 | 0.000185 |
| **Adjusted R-squared** | 0.8076 | | |
| **p-value** | < 2.2e−16 | | |

**Supplementary Table 3 – Statistical output for mixture models relating to plasma IgA**

| Model (Scheffé Polynomials) | Akaike Information Criterion | Degrees of freedom |
|---|---|---|
| 1 | 1160.604 | 4 |
| 2 | 1142.835 | 7 |
| 3 | 1149.336 | 11 |
| 4 | 1144.762 | 8 |
| Null | 1171.433 | 2 |

| **Model 2 Coefficients** | | | |
|---|---|---|---|
| **Components** | **Estimate (Std. Error)** | **t value** | **P(>\|t\|)** |
| Protein | 1244.4 (600.3) | 2.073 | 0.041706 |
| Fat | 2042.4 (371.3) | 5.501 | 5.31e−07 |
| Carbohydrate | 2110.2 (375.5) | 5.619 | 3.30e−07 |
| Protein:Fat | −3432.3 (1666.8) | −2.059 | 0.043043 |
| Protein:Carbohydrate | −4738.6 (1667.2) | −2.842 | 0.005804 |
| Fat:Carbohydrate | −6169.1 (1657.3) | −3.722 | 0.000385 |
| **Adjusted R-squared** | 0.7243 | | |
| **p-value** | < 2.2e−16 | | |

**Supplementary Table 4 – Statistical output for mixture models relating to plasma IgM**

| Model (Scheffé Polynomials) | Akaike Information Criterion | Degrees of freedom |
|---|---|---|
| 1 | 696.561 | 4 |
| 2 | 700.119 | 7 |
| 3 | 701.771 | 11 |
| 4 | 700.027 | 8 |
| Null | 693.555 | 2 |

**Supplementary Table 5 – 16S rRNA sequencing quality control**

| | Total reads | Reads post filtering | Reads post merging and chimera filtering | Reads retained |
|---|---|---|---|---|
| HP 1 | 135805 | 119366 | 97727 | 71.96% |
| HP 2 | 49009 | 43630 | 43301 | 88.35% |
| HP 3 | 120743 | 107426 | 91056 | 75.41% |
| HP 4 | 59370 | 51662 | 50927 | 85.78% |
| HP 5 | 71171 | 62355 | 61855 | 86.91% |
| HP 6 | 55324 | 49431 | 49288 | 89.09% |
| HP 7 | 24290 | 21876 | 21826 | 89.86% |
| HP 8 | 111569 | 96616 | 95494 | 85.59% |
| HC 1 | 89660 | 79158 | 78435 | 87.48% |
| HC 2 | 133887 | 119303 | 102883 | 76.84% |
| HC 3 | 52702 | 47334 | 46964 | 89.11% |
| HC 4 | 29954 | 26904 | 26437 | 88.26% |
| HC 5 | 121070 | 100002 | 94779 | 78.28% |
| HC 6 | 86078 | 71337 | 71144 | 82.65% |
| HC 7 | 69380 | 59366 | 59010 | 85.05% |
| HC 8 | 136477 | 117920 | 101334 | 74.25% |
| HF 1 | 44447 | 41330 | 40566 | 91.27% |
| HF 2 | 135418 | 126495 | 117867 | 87.04% |
| HF 3 | 13618 | 12835 | 12692 | 93.20% |
| HF 4 | 19284 | 18178 | 17971 | 93.19% |
| HF 5 | 23368 | 22069 | 21948 | 93.92% |
| HF 6 | 19495 | 18154 | 18134 | 93.02% |
| HF 7 | 120323 | 113333 | 103573 | 86.08% |
| HF 8 | 102020 | 95906 | 84963 | 83.28% |

**Supplementary Table 6 – Antibodies used for flow cytometry**

| Target | Conjugate | Clone | Manufacturer |
|---|---|---|---|
| CD45 | BV785 | 30-F11 | BioLegend |
| CD95 | APC | SA367H8 | BioLegend |
| GL-7 | FITC | GL7 | BioLegend |
| B220 | VioGreen | REA755 | Miltenyi Biotec |
| IgA | PE | 11-44-2 | eBioscience |

**Supplementary Table 7 – Primers used in this study and their sequence**

Pre-designed and validated primers used in this study. All primers were purchased from Sigma-Aldrich (KiCqStart SYBR® Green primers).

| **Gene target** | **Specie** | **Forward (5'-3')** | **Reverse (5'-3')** |
|---|---|---|---|
| *Pigr* | Mouse | AAGAACTCCAGAGATTTGGG | GTGGTAGTCACGATTTCATC |
| *Ccl28* | Mouse | GAGGTGTCTCATCATGTTTC | ATACGTTTTCTCTGCCATTC |
| *Tnfsf13* | Mouse | TCTATAGTCAGGTCCTGTTTC | GGCATACTTCTGATACATCG |
| *Baff* | Mouse | ATCTACAGCCAGGTTCTATAC | AGCTGAATCTCATCTCCTTC |
| *Tgfb* | Mouse | GGATACCAACTATTGCTTCAG | TGTCCAGGCTCCAAATATAG |
| *Tslp* | Mouse | CCTGAAACTGAGAGAAATGAC | ACACCCTTAGTATTCTGTCC |
| *Il10* | Mouse | AAGGGTTACTTGGGTTGCCA | AAATCGATGACAGCGCCTCAG |
| *Il4* | Mouse | CTGGATTCATCGATAAGCTG | TTTGCATGATGCTCTTTAGG |
| *Rpl13a* | Mouse | ATCCCTCCACCCTATGACAA | GCCCCAGGTAAGCAAACTT |
| *GAPDH* | Human | GAAGGTGAAGGTCGGAGTCA | CAGAGTTAAAAGCAGCCCTGG |
| *CCL28* | Human | GAGGTGTCTCATCATGTTTC | ATACGTTTTCTCTGCCATTC |
| *APRIL* | Human | CAGGTGTCTTCCATTTACAC | TGGAGAGAGGTTAAGTTTCG |
| *PIGR* | Human | GACCGAGTTTCAATCAGAAG | TTGTCATTGGCTCCAAATTC |

**Supplementary Figure 1**

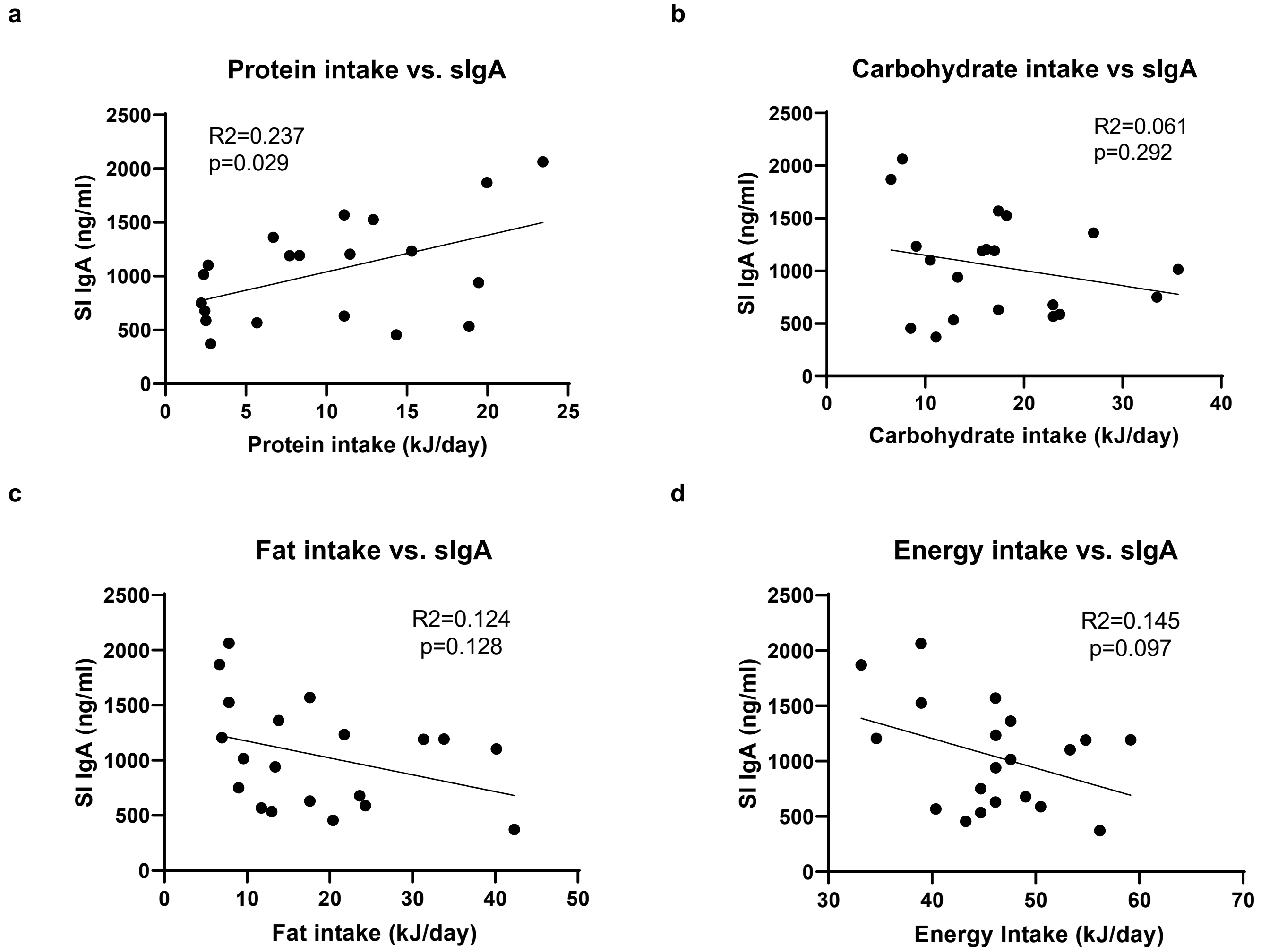

Supplementary Figure 1: Mice were fed on one of 10 diets encompassing a macronutrient range of 5-60% protein, 20-75% carbohydrate, and 20-75% fat and average food intake as well as sIgA quantified. Linear regression of (a) protein eaten, (b) carbohydrate eaten, (c) fat eaten or (d) total energy intake vs. sIgA levels were performed. Each dot represents the average intake versus the average quantified sIgA of a single cage (n=4 animals per cage). Macronutrient eaten and total energy intake is represented as kJ/day.

**Supplementary Figure 2**

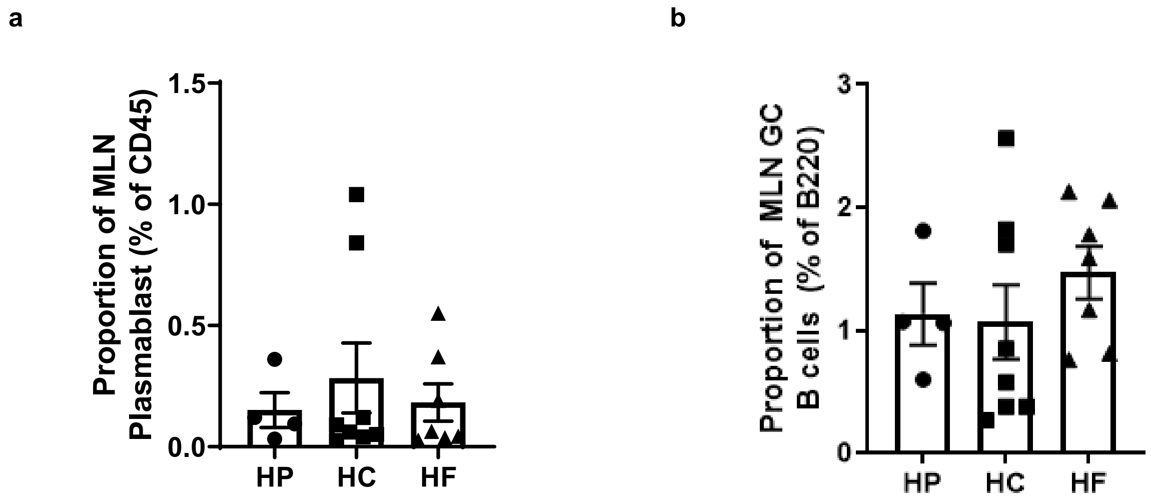

Supplementary Figure 2: (a) Proportion of B220$^{+}$IgA$^{+}$ plasmablast and (b) CD95$^{+}$GL7$^{+}$ germinal centre B cells in the mesenteric lymph nodes (n=4-8 per diet) were determined by flow cytometry from mice fed on either a HP, HC, or HF diet for 5 weeks. Data are represented as mean ± SEM.

**Supplementary Figure 3**

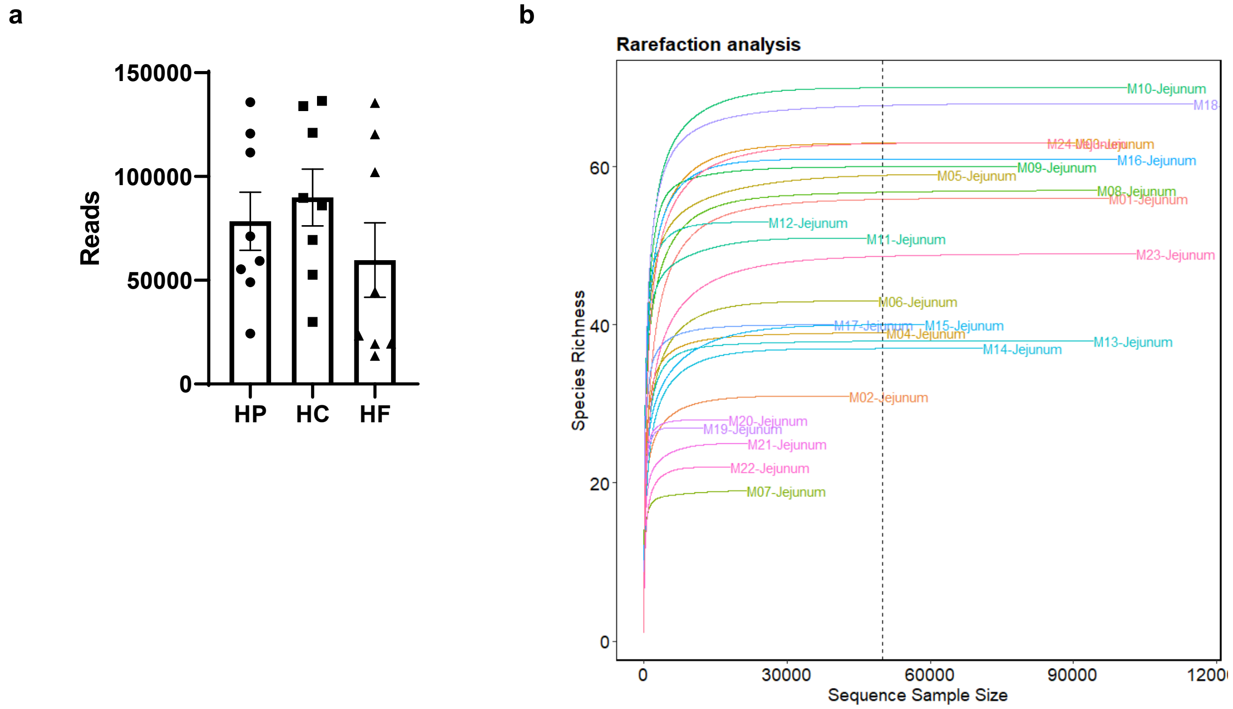

Supplementary Figure 3: DNA from luminal content was extracted for 16S rRNA DNA sequencing from mice fed on a high-protein, high-carbohydrate or high-fat diet for 5 weeks. (a) The total number of reads per samples were identified and no statistical significance between groups were found (Kruskal-Wallis test). (b) Rarefaction analysis was performed. Data are represented as mean ± SEM.

**Supplementary Figure 4**

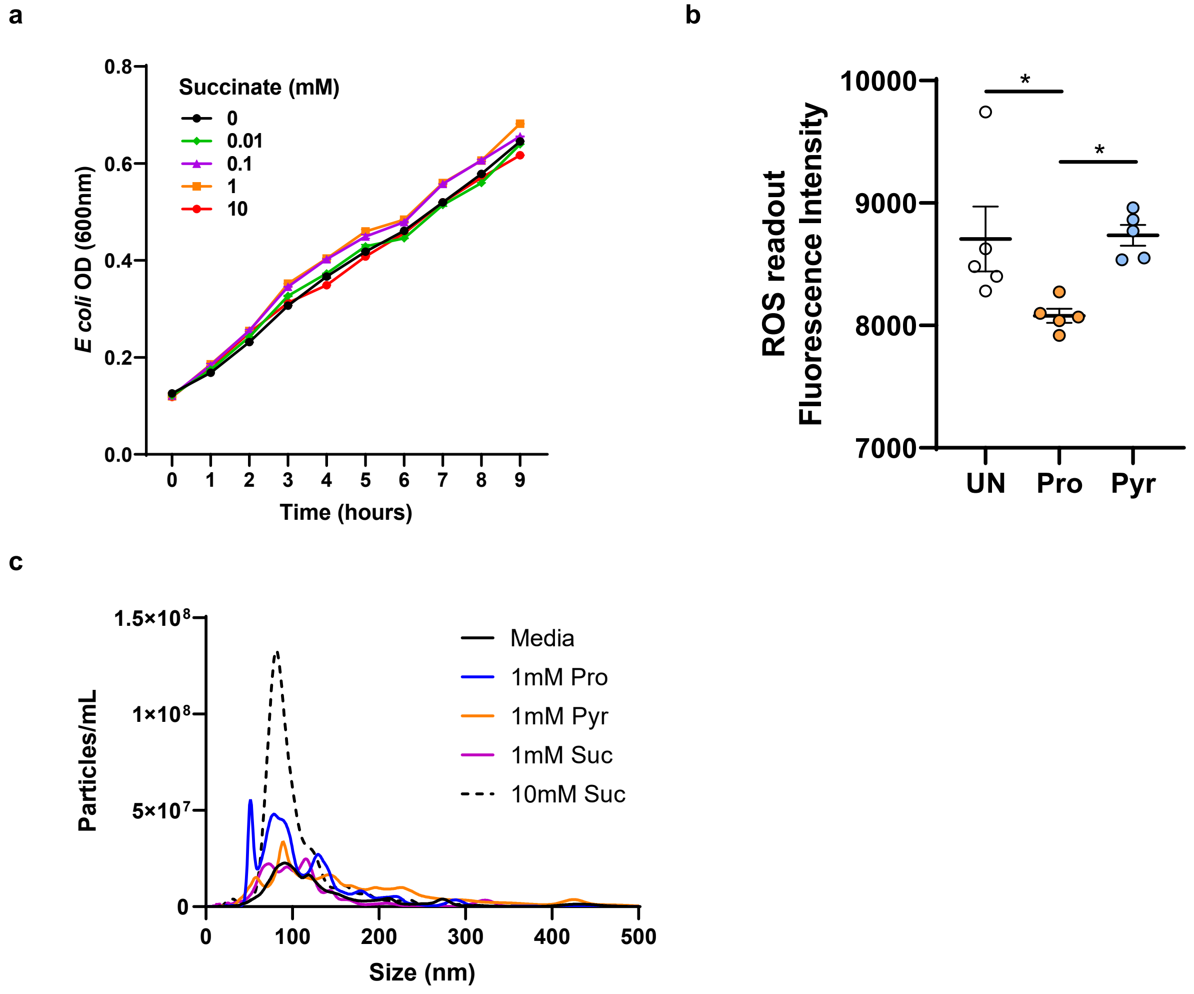

Supplementary Figure 4: (a) *E coli* was grown in LB broth in the presence of varying concentration of succinate and growth quantified by spectrophotometry (OD 600nm). (b) *E coli* was grown in LB broth in the presence of varying concentration of propionate (Pro) or pyruvate (Pyr) for 2 hours and ROS production quantified by the conversion of 2',7'-dichlorofluorescein diacetate to 2',7'-dichlorofluorescein (n=5 per condition). (c) *E coli* was grown in the presence of propionate (Pro), pyruvate (Pyr) or succinate (Suc) for 16 hours and extracellular vesicle was isolated and then quantified by nanoparticle tracking analysis (n=5-6 per condition) Data are represented as mean ± SEM. *$p < 0.05$
